# The DNA loop release factor WAPL suppresses Epstein-Barr virus latent membrane protein expression to maintain the highly restricted latency I program

**DOI:** 10.1101/2024.05.09.593401

**Authors:** Laura A. Murray-Nerger, Davide Maestri, Xiang Liu, Zhixuan Li, Italo Tempera, Mingxiang Teng, Benjamin E. Gewurz

## Abstract

Epstein-Barr virus (EBV) uses latency programs to colonize the memory B-cell reservoir, and each program is associated with human malignancies. However, knowledge remains incomplete of epigenetic mechanisms that maintain the highly restricted latency I program, present in memory and Burkitt lymphoma cells, in which EBNA1 is the only EBV-encoded protein expressed. Given increasing appreciation that higher order chromatin architecture is an important determinant of viral and host gene expression, we investigated roles of Wings Apart-Like Protein Homolog (WAPL), a host factor that unloads cohesins to control DNA loop size and that was discovered as an EBNA2-associated protein. WAPL knockout (KO) in Burkitt cells de-repressed LMP1 and LMP2A expression but not other EBV oncogenes to yield a viral program reminiscent of EBV latency II, which is rarely observed in B-cells. WAPL KO also increased LMP1/2A levels in latency III lymphoblastoid cells. WAPL KO altered EBV genome architecture, triggering formation of DNA loops between the LMP promoter region and the EBV origins of lytic replication (oriLyt). Hi-C analysis further demonstrated that WAPL KO reprograms EBV genomic DNA looping. LMP1 and LMP2A de-repression correlated with decreased histone repressive marks at their promoters. We propose that EBV coopts WAPL to negatively regulate latent membrane protein expression to maintain Burkitt latency I.

**Author Summary:** EBV is a highly prevalent herpesvirus etiologically linked to multiple lymphomas, gastric and nasopharyngeal carcinomas, and multiple sclerosis. EBV persists in the human host in B-cells that express a series of latency programs, each of which is observed in a distinct type of human lymphoma. The most restricted form of EBV latency, called latency I, is observed in memory cells and in most Burkitt lymphomas. In this state, EBNA1 is the only EBV-encoded protein expressed to facilitate infected cell immunoevasion. However, epigenetic mechanisms that repress expression of the other eight EBV-encoded latency proteins remain to be fully elucidated. We hypothesized that the host factor WAPL might have a role in restriction of EBV genes, as it is a major regulator of long-range DNA interactions by negatively regulating cohesin proteins that stabilize DNA loops, and WAPL was found in a yeast 2-hybrid screen for EBNA2-interacting host factors. Using CRISPR together with Hi-ChIP and Hi-C DNA architecture analyses, we uncovered WAPL roles in suppressing expression of LMP1 and LMP2A, which mimic signaling by CD40 and B-cell immunoglobulin receptors, respectively. These proteins are expressed together with EBNA1 in the latency II program. We demonstrate that WAPL KO changes EBV genomic architecture, including allowing the formation of DNA loops between the oriLyt enhancers and the LMP promoter regions. Collectively, our study suggests that WAPL reinforces Burkitt latency I by preventing the formation of DNA loops that may instead support the latency II program.

## Introduction

Epstein-Barr virus (EBV) infects >95% of adults and causes ∼200,000 cancers/year, including Burkitt and Hodgkin lymphomas and nasopharyngeal and gastric carcinomas [1–5]. Upon infection, the double-stranded DNA EBV genome is circularized and chromatinized, though much remains to be learned about how it folds into higher order structures. Upon B-cell infection, EBV switches between the pre-latency latency IIb and latency III programs [6–8], the latter of which expresses six Epstein-Barr nuclear antigens (EBNA) and two latent membrane proteins (LMP), LMP1 and LMP2A. LMP1 mimics signaling from activated CD40 receptors [9,10], whereas LMP2A subverts B-cell receptor signaling [11].

Microenvironmental cues trigger EBV to switch to latency IIa, where the Q promoter (Qp) and LMP promoters (LMPp) drive expression of EBNA1, LMP1, and LMP2A, respectively. Cytokines, in particular IL-15 and IL-21, downmodulate EBNA expression while supporting LMP1 expression [12–15]. Latency IIa B-cells further differentiate into memory cells, the EBV reservoir, where EBNA1 is the only viral protein expressed [1]. Latency IIa is observed in Hodgkin Reed-Sternberg tumor cells [1,2,16], while Burkitt lymphoma and gastric carcinoma use latency I [17] (**Fig. 1B**). However, much remains to be learned about the transition from latency IIa to latency I and about chromatin-based mechanisms that maintain latency I.

**Figure 1.**
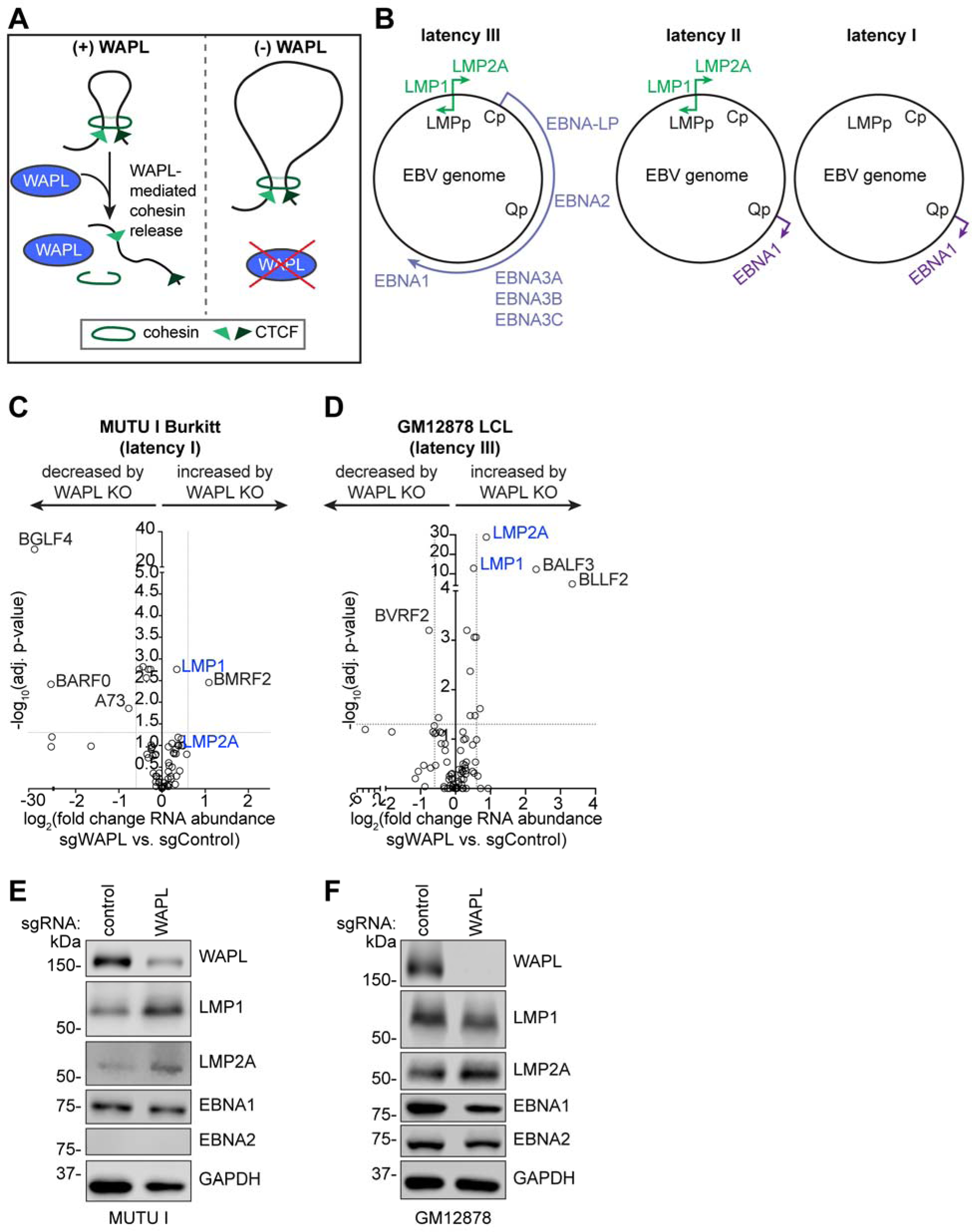
WAPL negatively regulates LMP1 and LMP2A expression. **(A)** Schematic of WAPL antagonism of cohesin-mediated DNA loop formation. WAPL releases cohesin to promote dissolution of chromatin loops. Upon WAPL KO, cohesin occupancy on chromatin increases, resulting in larger DNA loops. **(B)** Schematic diagram of EBV latency programs. **(C-D)** Volcano plots of RNA-seq analysis visualizing -log_10_(p-value) vs. log_2_(fold change of EBV mRNA abundance) from (C) Cas9+ MUTU I Burkitt lymphoma cells and (D) Cas9+ GM12878 LCLs expressing WAPL vs. control sgRNAs, from n = 3 independent biological replicates. **(E-F)** Immunoblot analysis of whole cell lysates (WCL) from (E) MUTU I cells and (F) GM12878 LCLs that expressed control or WAPL sgRNAs, as indicated, representative of n = 3 biological replicates.

Three-dimensional genome architecture is a major determinant of EBV gene expression [18–21]. The cohesin complex, comprised of SMC1, SMC3, and RAD21 subunits, forms a ring-shaped structure that encircles DNA to mediate long-range genomic interactions [22]. CTCF and cohesin are loaded onto discrete EBV and host genomic sites [18,21,23–31]. For instance, DNA loops juxtapose the EBV genomic origin of plasmid replication (OriP) enhancer with Cp and also with the LMP1/2p region to support latency III [24,31,32]. However, the OriP/LMPp loop is observed in latency I cells and is not sufficient to drive LMP1/2A expression [31].

Several factors limit DNA loop size [21,24–26]. First, paired CTCF sites block cohesin-driven loop extrusion to anchor DNA loops. Second, WAPL (wings apart-like protein homolog) [33,34] limits DNA loop size by opening a gate from which DNA can exit cohesin loops [35,36]. Consequently, large DNA loops are observed in WAPL deficient cells [34] (**Fig. 1A**). Notably, WAPL was discovered in a yeast-2 hybrid screen for host factors that associate with EBNA2 and was therefore originally named friend-of-EBNA2 (FOE) [37]. Despite this intriguing connection to EBV latency, WAPL roles in EBV-infected cells are unstudied.

Here, we tested the hypothesis that EBV utilizes WAPL to regulate viral gene expression. WAPL knockout (KO) in Burkitt cells de-repressed LMP1 and LMP2A, but not other EBV latency genes, suggestive of a switch to latency IIa. Long-range DNA analyses demonstrated that WAPL KO altered specific EBV genomic DNA loops, in particular at the LMP promoter regions and at the EBV oriLyt enhancers.

## Results

### WAPL is necessary for maintenance of EBV latency I

To test the role of WAPL in regulation of EBV gene expression, we knocked out WAPL in latency I Burkitt MUTU I or in latency III GM12878 lymphoblastoid cells (LCL) (**Fig. S1A, B**). WAPL KO did not significantly alter proliferation of either MUTU I or GM12878, even though it dramatically altered nuclear morphology (**Fig. S1A-D**), consistent with prior studies in EBV-negative cancer cell models [33,34].

To define how WAPL KO affects host and EBV gene expression, we performed RNA sequencing (RNA-seq) following acute WAPL KO or in control MUTU I and GM12878. While the expression of most EBV genes was not significantly changed by WAPL KO, LMP1 and LMP2A levels were significantly increased in MUTU I (**Fig. 1C, Table S1**). By contrast, EBNA2 was not substantially increased, suggesting an alternative mechanism increased LMP1/2A co-expression, perhaps reminiscent of latency II. Likewise, WAPL KO did not increase most EBV lytic genes or change EBV genome copy number (**Fig. 1C, 1E, S1E, Table S1**). WAPL KO also increased expression of LMP1/2A, but not of EBNA2 in GM12878 (**Fig. 1D, 1F, S1F, Table S1**).

We next interrogated WAPL KO effects on host gene expression. Consistent with LMP1 de-repression, LMP1/NF-κB target genes were amongst the most highly induced by WAPL [38], including mRNAs encoding the chemokines CCL3, CCL4 and CCL22, BIRC3 (which encodes cIAP2), and BCL2A1 (which encodes BFL1) (**Fig. S2A**). Gene ontology analyses identified that chemotaxis/chemokine pathways were the most highly upregulated by Burkitt WAPL KO (**Fig. S2B**). GM12878 WAPL KO also upregulated CCL3 and CCL4, together with antiviral responses and response to type II interferon (**Fig. S2C-D**).

### Subcellular distribution of de-repressed LMP1 and LMP2A

LMP1 and LMP2A signal from plasma membrane and endosomal sites, where they form puncta or membrane caps [39–44]. We asked whether WAPL KO induced typical LMP1 and LMP2A subcellular distribution. LMP1 puncta were observed in a significant proportion of WAPL KO, but rarely in control MUTU I (**Fig. 2A-B**). Similar results were obtained for LMP2A, in which LMP2A was de-repressed by WAPL KO and had similar subcellular distribution as in GM12878 (**Fig. 2C-D**).

**Figure 2.**
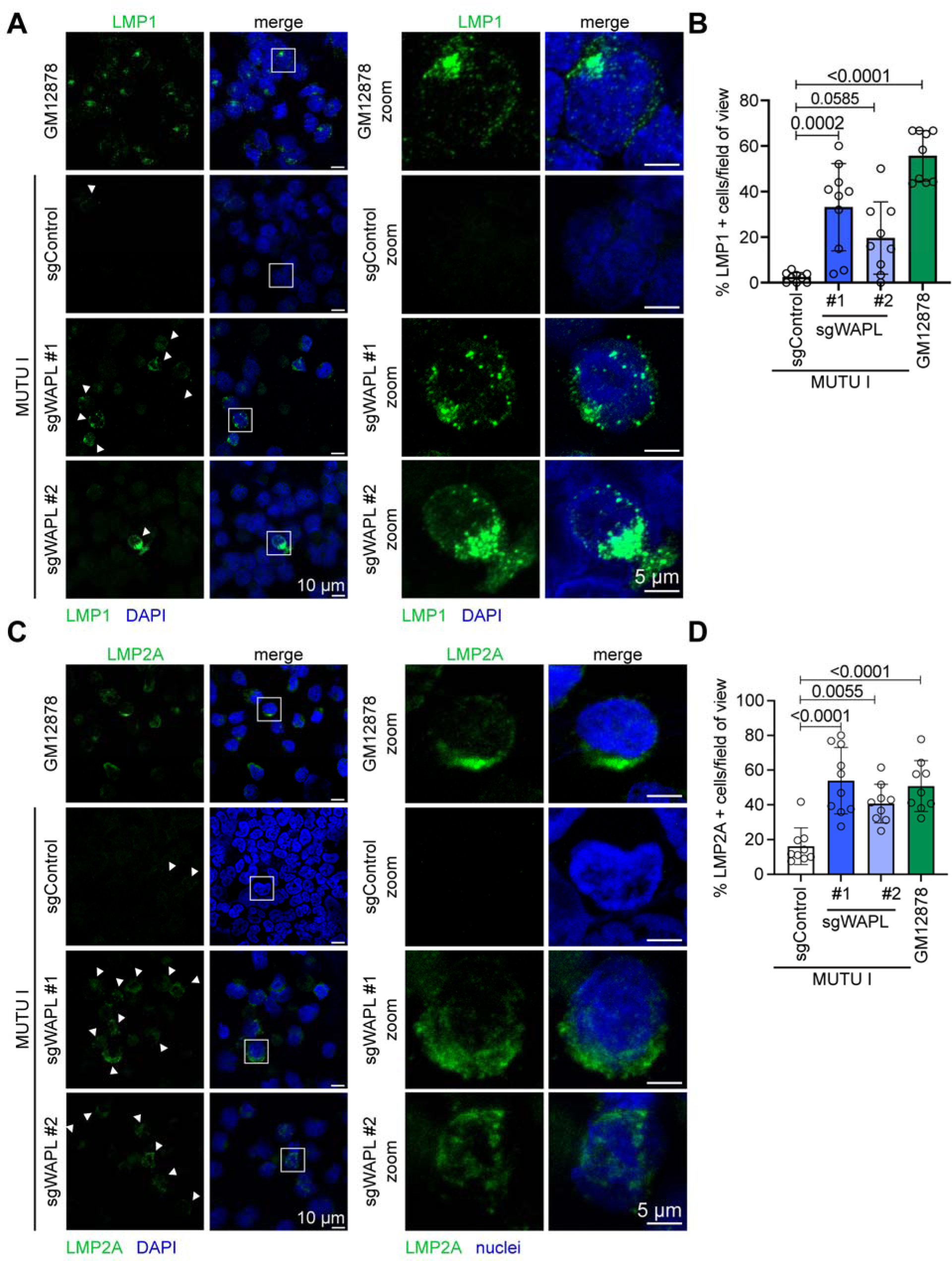
Subcellular distribution of LMP1 and LMP2A de-repressed by WAPL KO. **(A)** Representative confocal microscopy images from n = 3 biological replicates of anti-LMP1 (green) vs. nuclear DAPI (blue) staining of Cas9+ MUTU I cells that expressed control or WAPL sgRNAs, as indicated. Shown at right are zoomed images of a representative cell (indicated by the white box). (B) Mean ± standard deviation (SD) percentage of LMP1+ cells per field of view, from n = 3 fields of view from each of three biological replicates. *P*-values shown as calculated by one-way ANOVA. (B) Representative confocal microscopy images from n = 3 biological replicates of anti-LMP2A (green) vs. nuclear DAPI (blue) staining of Cas9+ MUTU I that expressed control or WAPL sgRNAs with zoomed images presented to the right, as in (A). (D) Mean ± SD percentage of LMP2A+ cells per field of view, from n = 3 fields of view from each of three biological replicates. *P*-values shown as calculated by one-way ANOVA.

Since latency IIa B cell models are unavailable, we next asked whether LMP1 and LMP2A formed membrane puncta in WAPL KO P3HR-1 Burkitt cells, which harbor an EBNA2 deletion [45–48]. Indeed, WAPL KO de-repressed LMP1 and LMP2A in P3HR-1, which formed characteristic puncta (**Fig. S3A-E**), indicating that WAPL is required to repress Burkitt LMP expression even in the absence of EBNA2. However, the percentage of cells that de-repressed LMP1 and LMP2A were somewhat lower than in MUTU I or GM12878. This may be related to disruption of EBV genomic architecture by the deletion present in P3HR-1.

### WAPL regulates LMP region looping

To test the hypothesis that WAPL KO altered EBV genomic architecture to de-repress LMP1/2A, we performed EBV genomic Hi-C, which measures long-range DNA contacts using proximity ligation with high-throughput sequencing [28,49,50] (**Fig. 3A**). At a cutoff of FDR < 0.05 and Z-score > 1, Hi-C identified that 60 EBV genomic loops were gained upon WAPL KO (**Fig. 3B, Table S2**), including between the LMP region and the rightward oriLyt (oriLyt^R^) enhancer. A loop was also gained between the LMP region and BKRF2, which in turn looped to the BLRF2 and EBNA-1 region (**Fig 3B**). WAPL depletion significantly decreased 138 EBV DNA loops at the cutoff of FDR < 0.05 and Z-score < -1 (**Fig. 3C, Table S2**), including from the LMP region to multiple EBV genomic locations, including the leftward oriLyt (oriLyt^L^) (**Fig. 3C**).

**Figure 3.**
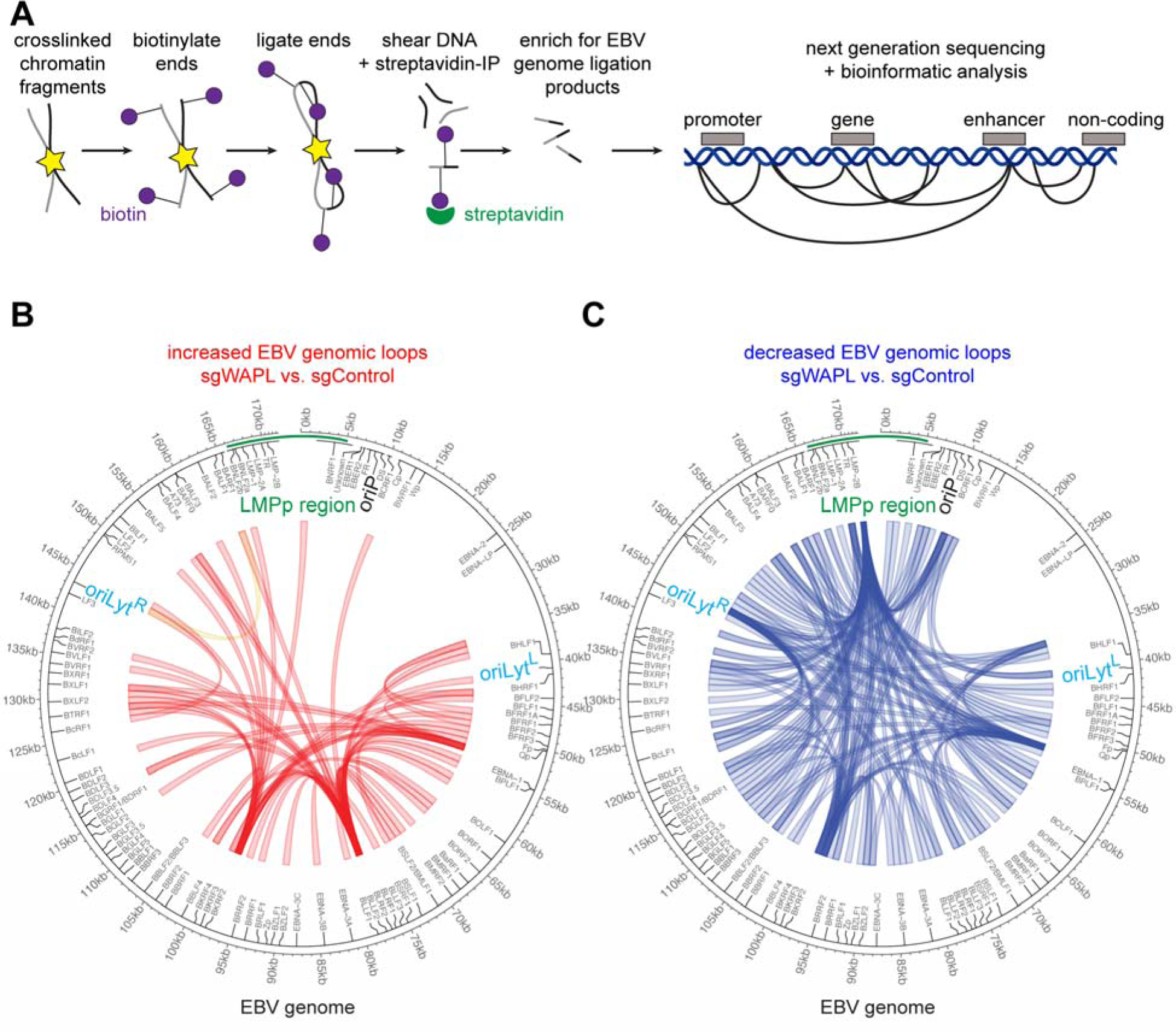
WAPL KO alters higher order latency I Burkitt EBV genome conformation. **(A)** Schematic of Hi-C workflow and output. Exposed DNA ends were biotinylated and then ligated to capture close DNA contacts. Ligated DNA was sheared, and biotinylated DNA was precipitated. EBV DNA was captured to enhance viral DNA Hi-C signal. **(B)** Hi-C maps of EBV genomic loops that were enriched in WAPL KO vs. control MUTU I cells, from n = 2 biological replicates. LMPp and oriLyt regions are indicated. **(C)** Hi-C maps of EBV genomic loops that were depleted in WAPL KO vs. control MUTU I cells, from n = 2 biological replicates, as in (B).

We next used HiChIP [51] to define how WAPL KO altered long-range EBV genomic interactions between areas of activated chromatin [52,53], marked by histone 3 lysine 27 acetyl (H3K27Ac) (**Fig. 4A**). HiChIP identified a higher frequency of interactions between *LMP* and both oriLyt regions (**Fig. 4B-D, Fig S4A-B, Table S3**). By contrast, WAPL KO decreased interactions between H3K27Ac-marked LMP and several other EBV genomic regions (**S4A-B**). Thus, both Hi-C and HiChIP detected formation of a loop between oriLyt^R^ and the LMP promoter region formed upon WAPL KO.

**Figure 4.**
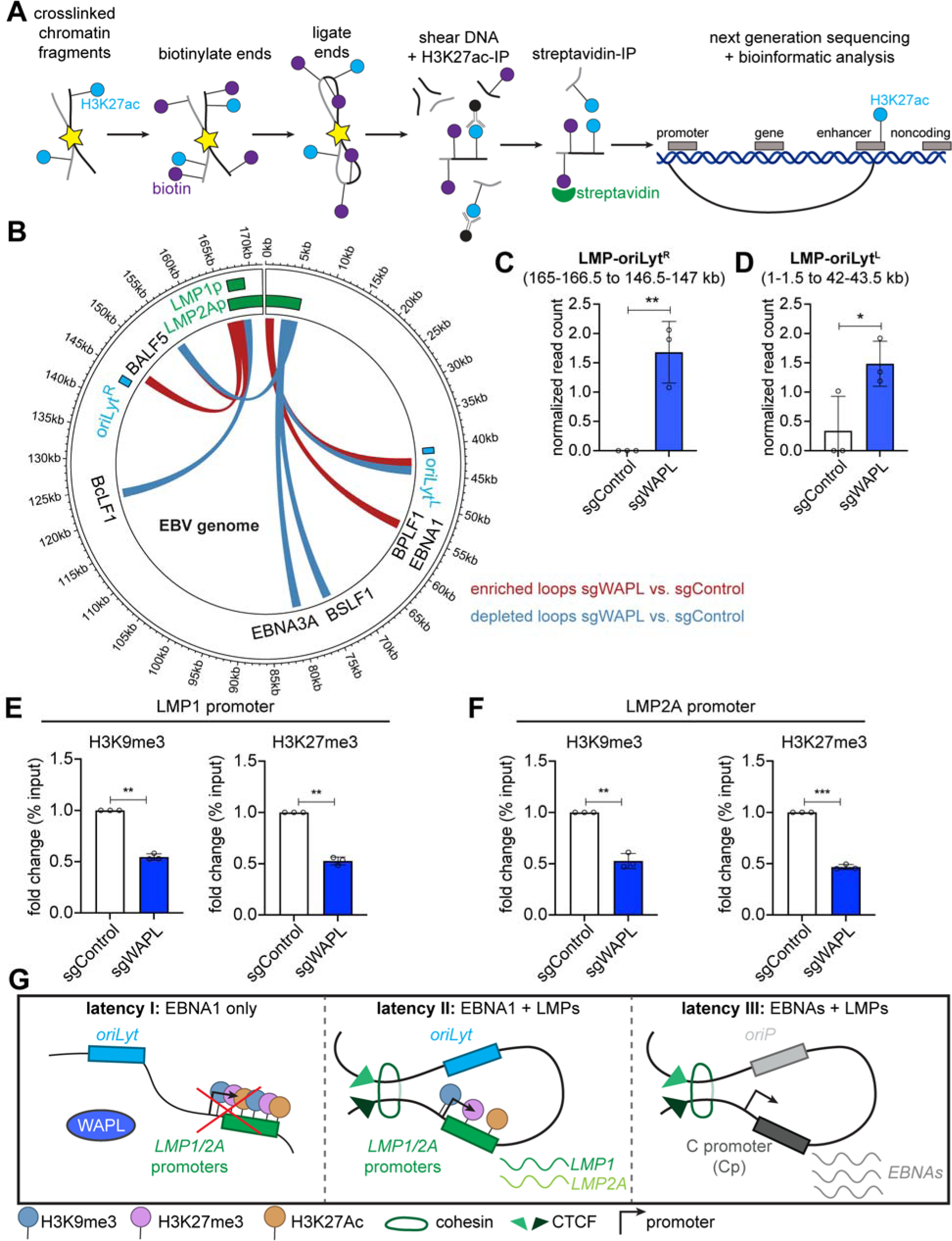
WAPL KO alters latency I Burkitt EBV genomic activated chromatin loops and represses LMP promoter epigenetic marks. **(A)** Schematic of H3K27Ac HiChIP sample preparation and output. Chromatin was formaldehyde crosslinked and fragmented. Exposed DNA ends were biotinylated and then ligated to capture close DNA contacts. Ligated DNA was sheared, DNA was immunopurified by α−H3K27Ac antibody, and biotinylated DNA was captured via streptavidin. **(B)** EBV genomic H3K27Ac HiChIP map depicting loops enriched (red) versus depleted (blue) in WAPL KO MUTU I cells, relative to levels in control cells, from n = 3 biological replicates. **(C-D)** Normalized (C) LMP region-oriLyt^L^ loop and (D) LMP region-oriLyt^R^ loop read counts from n = 3 replicates, as in (B). EBV genome kilobase coordinates for each looping region are indicated at top. * *P* ≤ 0.05, ** *P* ≤ 0.01, as calculated by a two-tailed Student’s t-test. **(E-F)** ChIP-qPCR analysis of H3K9me3 and H3K27me3 abundances at the (E) LMP1 promoter and (F) LMP2A promoter in Cas9+ MUTU I cells expressing control or WAPL sgRNAs. Shown are mean fold change of the percentage input values ± SD from n = 3 biological replicates. ** *P* ≤ 0.01, *** *P* ≤ 0.001, as calculated by a two-tailed Welch’s t-test. **(G)** Model of WAPL effects on EBV genomic architecture. When present, WAPL releases cohesin at the targeted DNA loop (latency I), which inhibits LMP expression. In the absence of WAPL antagonism, cohesins are loaded onto the EBV genome to form loops between the LMP promoter region and oriLyt regions. Juxtaposition of the oriLyt enhancer reduces repressive H3K9me3 and H3K27me3 marks and supports *LMP1* and *LMP2A* co-expression in the absence of EBNA2 (latency II). In latency III, an alternative loop forms between the oriP and the Cp to drive expression of all of the EBNA genes.

We next characterized how WAPL KO altered LMP1 promoter region histone marks. WAPL KO significantly increased repressive histone 3 lysine 9 and lysine 27 trimethylation (H3K9me3/H3K27me3) levels at both the LMP1 and LMP2A promoter regions (**Fig. 4E-F**).

While polycomb repressive complex I mediated histone 2A lysine 119 monoubiquitination (H2AK199Ub) represses Burkitt LMP1 and LMP2A [54], its levels were not significantly changed by WAPL KO at LMP1 or LMP2A promoter regions (**Fig. S5A-B**). WAPL KO did not significantly change H3K27Ac marks at the LMP1 promoter and decreased them at the LMP2A promoter (**Fig. S5A-B**). These results suggest that WAPL supports EBV latency I by altering EBV genomic structure to increase repressive LMPp H3K9me3 and H3K27me3 marks to enforce latency I maintenance (**Fig 4G**).

## Discussion

Much remains to be learned about epigenetic mechanisms that maintain latency I. Here, we found that the cohesin release factor WAPL suppresses LMP1 and LMP2A expression in Burkitt latency I by supporting higher order EBV genomic architecture. WAPL KO triggered DNA loops between *oriLyt* and LMPp, decreased LMPp repressive H3K9me3/H3K27me3 marks, and de-repressed LMP1/2A co-expression, even in the absence of EBNA2. These results highlight an important WAPL role in preventing reversion to latency II.

Loss of WAPL permits cohesin to slide beyond host CTCF anchors and enlarges host DNA loops [33]. Our findings suggest that WAPL KO likewise regulates EBV genome architecture. To our knowledge, WAPL effects on viral genomes have not previously been defined. Furthermore, our results suggest that EBV genomic structure may be distinct between germinal center B-cells in latency IIa versus memory B-cells in latency I. Therefore, important future objectives will be to determine (1) whether WAPL abundance or activity differs between EBV-infected germinal center and memory B-cells and (2) to define germinal center versus memory B-cell EBV genomic architecture as technologies become available to do so on the single cell level as these populations are rare *in vivo*.

WAPL KO reduced LMPp histone repressive marks in latency I, suggesting that WAPL supports an EBV genomic configuration that contributes to LMP1 and LMP2A repression. While we cannot rule out that WAPL KO instead alters a host factor that alters LMPp epigenetic marks, RNAseq analysis did not reveal significant changes in the expression of H3K9me3 or H3K27me3 writers or erasers. Thus, we instead favor the model that WAPL prevents the formation of loops between oriLyt and LMPp that induce LMP1/2A co-expression. Notably, DNA loops between oriLyt and LMP promoter regions have been described in gastric carcinoma and natural killer cells [26,55], but not previously in B-cells. Instead, in latency III, cohesin and CTCF bind to the LMP1 and LMP2A control region at a site that overlaps the first *LMP2A* intron and the LMP1 3′ untranslated region to drive a loop between the oriP enhancer and LMPp in support of LMP1 and LMP2A expression. However, the oriP:LMPp loop is present in MUTU I, where LMP1/2A are epigenetically silenced [31], suggesting that additional mechanisms repress LMP expression in latency I. Furthermore, cohesin knockdown elevates LCL LMP1/2A levels, and deletion of the LMP region CTCF site increases repressive LMP2p H3K9me3 and DNA methylation marks [24,31,32], consistent with our finding that DNA loop(s) can repress LMP1/2A.

Although WAPL was discovered as an EBNA2 binding partner [37], the role of WAPL in EBV genome regulation had remained unstudied. Since EBNA2 is a major inducer of LMP1 and LMP2A in EBV latency III, an intriguing possibility is that EBNA2 not only activates LMPp chromatin but may also dismiss WAPL from this key EBV genomic region. In this manner, EBNA2 may alter EBV genomic architecture to reduce H3K9me3/H3K27me3 repressive marks in support of LMP expression in newly infected cells. It may also work in latency III in a similar manner while being supported by recruitment of co-activators and effects on DNA hypomethylation [56,57]. Taken together, we now propose that WAPL prevents loops between oriLyt and LMPp to repress LMP1/2A in latency I, whereas a distinct oriP/LMPp loop supports LMP expression in latency III.

In conclusion, EBV coopts WAPL in latency I to regulate higher order EBV genome architecture to restrict LMP1 and LMP2A expression. It provides a new latency II B-cell model and lays the foundation for future studies of how WAPL remodels enhancer/promoter communication for EBV and for the three-dimensional genome regulation of other double stranded DNA viruses.

## Materials and Methods

### RNA-seq

RNA was extracted from B-cells and poly-A enrichment was performed prior to library preparation and next generation sequencing. Reads were mapped to the hg19 human (GRCh37) and Akata EBV genomes. Salmon (v1.0.0) was used to quantify the transcripts [58], and DESeq v1.14.1[59] was used to determine differentially expressed genes. Genes that had a log_2_(fold change) of at least 0.6 (actual fold change of 1.5) and an adjusted p-value of < 0.05 were considered significant.

### Hi-C

The Hi-C assay was performed as previously described [28]. Significantly changed associations (FDR < 0.05 and Z-score > 1 or < -1) were plotted as circos graphs using the circlize package (version 0.4.12) of R (version 4.0.5) [60].

### HiChIP

HiChIP was performed as previously described [51]. In brief, HiChIP read loops between EBV genomic bins (1.5kb) were quantified followed by normalization using loops per 10k total read pairs. Wilcoxon Rank Sum test was used to evaluate loop differences between conditions. Top differential loops (p-value < 0.1, difference > 3 normalized read pairs, mean read pairs ≥ 2 in at least one condition) were visualized by circlize v0.4.15 R package [60].

## Data availability

Extended methods are available in the supplementary information. RNA-seq, HiChIP, and Hi-C data are deposited on the NIH GEO database using accession numbers GSE248336, GSE248335, and GSE264502 respectively. All figures were made using commercially available GraphPad, Adobe Illustrator, or packages in R.

## Acknowledgements

We thank members of the Gewurz and Zhao labs for helpful feedback and Rui Guo for advice on RNA-seq data analysis. This work was supported by T32 AI007245 and F32 AI172329 to L.A.M.N., by U01 CA275301, R01 CA228700, R01 AI164709, and R21 AI170751 to B.E.G., and by P01 CA269043 to I.T. and B.E.G. I.T. and D.M. were also supported by R01 AI130209. We appreciate the support of the Molecular Biology Genomics Core at the Dana Farber Cancer Center for RNA-seq and Hi-ChIP data acquisition.

## Author contributions

L.A.M.N., D.M, Z.L., and B.E.G. designed experiments. L.A.M.N., D.M., and Z.L. performed experiments. L.A.M.N., D.M., X.L., Z.L., I.T., M.T., and B.E.G. analyzed data. L.A.M.N., D.M., X.L., Z.L., I.T., M.T., and B.E.G. wrote the manuscript.

## Competing interests

The authors have no conflicts of interest.

## Notes

### Competing Interest Statement

The authors have declared no competing interest.

