## Supplementary Information for "The DNA loop release factor WAPL suppresses Epstein-Barr virus latent membrane protein expression to maintain the highly restricted latency I program"

Murray-Nerger, et. al.

**Extended methods**

**Cell lines and culture**

The B-cell lines used in this study are from the following sources: P3HR-1 ZHT/RHT Cas9+ (original cell line gift from Dr. Eric Johannsen, Cas9+ cell line from [1]), MUTU I Cas9+ (original cell line gift from Dr. Jeff Sample, Cas9+ cell line generated in our lab), and GM12878 Cas9+ (original cell line from Coriell, Cas9+ made in our lab). B-cells were cultured in Roswell Park Memorial Institute 1640 Medium (RPMI) (Life Technologies) supplemented with 10% fetal bovine serum (FBS) (v/v) (Thermo Fisher) and 1% penicillin-streptomycin (P/S) (v/v) (Life Technologies). 293T cells were purchased from ATCC and cultured in Dulbecco's Modified Eagle Medium (DMEM) (Thermo Fisher Scientific) supplemented with 10% FBS and 1% P/S. Cells were grown at 37 °C and 5% $CO_2$ and were tested for mycoplasma using a MycoAlert Mycoplasma Detection Kit (Lonza).

**CRISPR-Cas9 editing**

Lentiviral sgRNA expression vectors were generated by cloning the sgRNA of interest into the pLenti-Guide-puro vector (Addgene #52963) by digesting the vector with BsmBI and ligating in the annealed sgRNA sequence. Plasmids containing the correct sequences were confirmed via Sanger sequencing. Guide sgRNA sequences used were the following: sgControl (TTGACCTTTACCGTCCCGCG), sgWAPL #1 (CGTTCCATAGTATCCTGTA), and sgWAPL #2 (TCAGAAATATCTCCAATCAA). 293T cells were transfected with VSV-G, psPAX2, and the pLenti-puro construct. Lentivirus was collected 48 and 72 h post-transfection and used to infect B-cells. Cells were selected for 6 days using puromycin (3 µg/mL), and WAPL KO was confirmed by immunoblot.

**Cell Titer Glow analysis**

CellTiter-Glo live cell number analysis (Promega) was performed as per the manufacturer's instructions. Cells were collected at the indicated time points and were washed once with 1x PBS. 30 µL washed cells were used per assay.

**RNA-seq library preparation and analysis**

RNA was extracted from 2 million cells in biological triplicate using a RNeasy extraction kit (Qiagen), and an on-column DNA digestion step was performed. Bioanalyzer analysis was used to confirm RNA quality (RIN score > 7). For library preparation, 1 µg of RNA was used for each sample, poly-A enrichment using the NEBNext Poly(A) mRNA Magnetic Isolation Module (#E7490, New England Biolabs) was performed, and libraries were constructed using NEBNext Ultra RNA Library Prep Kit for Illumina (#E7760L, New England Biolabs) as per the manufacturer's instructions. Library quality was confirmed via Bioanalyzer analysis. Libraries were multi-indexed, pooled, and sequenced on an Illumina NextSeq 500 sequencer using single-end 75-bp reads (Illumina) at the Molecular Biology Genomics Core Facility at the Dana-Farber Cancer Institute. Single-end reads were mapped to the hg19 human (GRCh37) and Akata EBV genomes. Salmon (v1.0.0) was used to quantify the transcripts [2], and DESeq v1.14.1[3] was used to determine differentially expressed genes. Genes that had a $\log_2$(fold change) of at least 0.6 (actual fold change of 1.5) and an adjusted p-value of < 0.05 were considered significant.

**Immunoblot analysis**

Samples were lysed in 1x Lammeli buffer by vortexing for 15 s and then either heating at 95°C for 10 min or heating at 70°C for 10 min (for LMP2A blotting). Equal amounts of sample were loaded onto a tris-glycine SDS-polyacrylamide gel. Proteins were transferred to a nitrocellulose membrane, which was cut and blocked with 5% milk in 1x TBS. All primary antibodies were diluted into 1x TBST (0.1% Tween-20) and incubated overnight at 4°C. The primary antibodies used are the following: WAPL (1:1000, rabbit, Cell Signaling D9J1U), LMP1 (1:2000, mouse, Abcam CS1-4 ab78113), LMP2A (1:500, rat, Abcam 15F9 ab59028), GAPDH (1:20,000, mouse, Millipore MAB374), EBNA2 (mouse, clone PE2, gift from Fred Wang). Membranes were washed with 1x TBST and incubated for 1 h at room temperature with the following secondary antibodies used at a 1:4000 dilution: anti-rabbit IgG, HRP-linked (Cell Signaling Technologies 7074V), anti-mouse IgG, HRP-linked (Cell Signaling Technology 7076V), and anti-rat IgG, HRP-linked (Cell Signaling Technology 7077S). Membranes were then washed with 1x TBST and incubated with SuperSignal™ West Pico PLUS Chemiluminescent Substrate (Thermo Fisher Scientific) for 2 min before imaging on a Li-Cor Odyssey Fc.

**Immunofluorescence microscopy**

Collected B-cells were washed with 1x PBS and dried on slides for 30 min at 37°C. Cells were fixed with 4% paraformaldehyde (10 min), permeabilized with 0.5% Triton-X100 (5 min), and blocked for 1 h with 1% BSA. Samples were incubated with primary antibody for 1 h at 37°C, washed with PBS, incubated with secondary antibody for 1 h at 37°C, washed with PBS, incubated with 1:5000 Hoechst (Thermo Fisher) for 15 min at 37°C, and then mounted using ProLong™ Diamond Antifade Mountant (Thermo Fisher). Images were collected using a 63x objective on a Zeiss LSM 800 microscope. Image analysis was performed in ImageJ by counting the number of nuclei and the number of cells expressing either LMP1 or LMP2A. Antibodies used are as follows: LMP1 (1:250, mouse, Abcam CS1-4 ab78113), LMP2A (1:250, rat, Abcam 15F9 ab59028), anti-Ms 488 (1:500, Goat anti-Rabbit IgG (H+L) Cross-Adsorbed Secondary Antibody, Alexa Fluor 488), anti-Rat 647 (1:500, Goat anti-Rabbit IgG (H+L) Cross-Adsorbed Secondary Antibody, Alexa Fluor 647).

**Hi-C sample preparation and analysis**

The Hi-C assay was performed as previously described [4]. Briefly, 5 million cells per condition were collected for in-situ Hi-C. Libraries of total ligation products were produced using Ultralow Library Systems V2 (Tecan Genomics, part no. 0344NB-32) as per the manufacturer's protocol. Purified libraries were then enriched for only EBV genome ligation products using myBaits enrichment kit as per the manufacturer's protocol. Libraries were sequenced using the Illumina HiSeq 2500 sequencing platform with paired-end 75bp read length. Hi-C data was preprocessed using HiC-Pro v2.10.0 pipeline [5] with default settings using the NC_007605.1 version of the EBV genome at 1kb resolution. Replicates per condition were then merged and HiCcompare software [6] was used to estimate significance of differential contact based on raw count matrix files. Samples were first normalized by library size and then by loess normalization. To determine filtering parameters the *filter_params()* function was used, applying numChanges = 1500, FC = 3 given the high resolution of the Hi-C matrices. A minimum average expression (A.min) was set to "15" and the p.method to "fdr" for the function *hic_compare()*. Significantly changed associations (FDR < 0.05 and Z-score > 1 or < -1) were plotted as circos graphs using the circlize package (version 0.4.12) of R (version 4.0.5) [7]. Only associations between regions that are 5kb distant were considered.

**HiChIP sample preparation**
HiChIP was performed as previously described in the Mumbach et al. methods paper [8]. Briefly, 10 million cells were collected and crosslinked with freshly made 1% formaldehyde, and crosslinking was quenched with 125 mM glycine. Crosslinked cells were lysed with ice-cold Hi-C lysis buffer at 4°C for 30 minutes with rotation. Pelleted nuclei were harvested and digested with 375U of MboI restriction enzyme (NEB, R0147) at 37°C for 2 hours with rotation. Next, DNA ends were filled in and marked with biotin-dATP (Thermo, 19524016). T4 DNA ligase (NEB, M0202) was used for proximity ligation at room temperature for 4 hours with rotation. After ligation, pelleted nuclei were lysed with ice-cold nuclear lysis buffer and sonicated with Covaris M220 for 240 seconds. ChIP dilution buffer was used to dilute each sample to achieve a SDS concentration of 0.1%. After preclearing with Protein A magnetic beads, 7.5 μg of anti-H3K27ac antibody (abcam, ab4729) were added for every 10 million cells and incubated overnight at 4°C. Protein A magnetic beads were used to capture DNA-protein complexes. Beads were washed three times each in Low Salt Wash Buffer (20 mM Tris pH 8, 2 mM EDTA, 150 mM NaCl, 1% Triton X-100, 0.1% SDS), High Salt Wash Buffer (20 mM Tris pH 8, 2 mM EDTA, 500 mM NaCl, 1% Triton X-100, 0.1% SDS), and LiCl Wash Buffer (10 mM Tris pH 8, 1 mM EDTA, 0.25 M LiCl, 1% NP40, 1% sodium deoxycholate), and DNA was eluted in Elution Buffer (50 mM NaHCO3, 1% SDS). Crosslinks were then reversed via Proteinase K digestion, and DNA was purified with the QIAquick PCR purification kit (Qiagen). Samples were prepared for sequencing using the Illumina Tagment DNA Enzyme and Buffer Kits (Illumina, 20034197) per the manufacturer's instructions. Samples were sequenced on an Illumina NextSeq 500 sequencer (PE 75bp) at the Molecular Biology Genomics Core Facility at the Dana-Farber Cancer Institute.

**HiChIP data analysis**
HiChIP data analysis of each sample was performed as previously described [9]. Briefly, reads of biological replicates were processed with the HiC-Pro v3.1.0 pipeline [5] against a MboI digested hg19 with EBV genome build (NCBI Refseq: NC_007605.1). For the human genome, Hichipper v0.7.7 [10] (parameter: EACH, ALL) was used to identify long-rang chromatin interactions based on valid HiChIP read pairs selected by HiC-Pro. Differential analysis of long-range interactions was performed by the diffloop v1.24.0 [11]. Loop biases were filtered based on the binomial model described by mango using *mangoCorrection* [12]. Loops with FDR < 0.01 and range > 5kb were kept for differential analysis. Differential loops between two conditions were estimated by considering reproducibility across replicate samples using *quickAssocVoom*. Significant differential loops (Benjamini-Hochberg FDR< 0.05) were selected for visualization. For the EBV genome, genomic bins of 1500 bps length were treated as potential loop anchors. All pair-wise combinations of bins were quantified for their strength of potential interactions using HiChIP read pairs supporting the interacting two bins. Only read pairs labeled as *allValidPairs* by HiC-Pro were selected for quantification. Neighbored bin pairs and bin pairs with zero quantification in all samples were removed. Read pair quantification was normalized based on read pairs per 10k per EBV genome. Wilcoxon Rank Sum test was used to evaluate loop differences between conditions. Top differential loops (p-value < 0.1, difference > 3 normalized read pairs, mean read pairs ≥ 2 in at least one condition) were visualized by circlize v0.4.15 R package [7].

**ChIP-qPCR**

Cells were harvested and cross-linked with 1% formaldehyde for 10 min at room temperature with rotation and crosslinking was quenched 0.3 M glycine for an additional 5 min. Cells were washed twice with cold PBS and stored at -80°C. Cells were lysed in lysis buffer (50 mM Tris pH 8, 10 mM EDTA, 1% SDS, protease inhibitor (Sigma, catalog #11873580001)) for 20 min on ice. Cell lysate was sonicated for 10 cycles (30s on/30s off) in a Diagenode water bath sonicator. Correct fragment size was confirmed via agarose gel electrophoresis. Samples were centrifuged at 13,000 rpm for 10 min at 4°C. Soluble chromatin was diluted 1:10 in ChIP dilution buffer (16.7 mM Tris pH 8, 1.2 mM EDTA, 167 mM NaCl, 1.1% TritonX-100, 0.01% SDS, and protease inhibitor). Chromatin from 1 million cells was combined with 2 µg antibody and 20 µL pre-washed magnetic protein A/G beads (Thermo) and incubated overnight with rotation at 4°C. The following day, a series of washes was performed as follows: twice with low salt buffer (20 mM Tris pH 8, 2 mM EDTA, 150 mM NaCl, 1% Triton X-100, 0.1% SDS), twice with high salt buffer (20 mM Tris pH 8, 2 mM EDTA, 500 mM NaCl, 1% Triton X-100, 0.1% SDS), once with LiCl buffer (10 mM Tris pH 8, 1 mM EDTA, 0.25 M LiCl, 1% NP40, 1% sodium deoxycholate), and once with TE (IDT). For elution and crosslink reversal, beads were resuspended in 100 µL elution buffer (100 mM NaHCO3, 1% SDS), supplemented with 5% v/v Proteinase K (NEB), and incubated at 65°C for two hours with shaking at 400 rpm. Input samples were also incubated with 5% v/v Proteinase K at the same time. The QIAquick PCR purification kit (Qiagen) was used to purify the eluted DNA. Quantification was performed via qPCR run on a Biorad CFX Connect Real Time System and normalized to the percentage of input DNA (% input). Antibodies used are as follows: H2AK119Ub (Cell Signaling, 8240S), H3K9me3 (Active Motif, 39161), H3K27me3 (Cell Signaling, 9733S), H3K27Ac (Abcam, ab4729), and anti-IgG (Cell Signaling 2729S). Primers used are as follows: LMP1p_F (5'-ACGTCAGAGTAACGCGTGTTTC-3'), LMP1p_R (5'-GCAGACCCCGCAAATCC-3'), LMP2Ap_F (5'-GCGCCGCGGTTTCAG-3'), and LMP2Ap_R (5'-TTACGCCCCAGCAAGCTT-3').

**Quantification of EBV genomes**

Cells were harvested and DNA was extracted using a Blood and Cell Culture DNA minikit (Qiagen). Extracted DNA was diluted to 10 ng/µL, and qPCR for the viral BALF5 gene was performed on a Biorad CFX Connect Real Time System. A standard curve was generated via serial dilution of the pHAGE-BALF5 plasmid (generated in our lab) starting at 25 ng/µL, and the resulting linear regression equation was used to quantify the number of copies of the viral genome. Primers used are as follows: BALF5_F (5'-GAGCGATCTTGGCAATCTCT-3') and BALF5_R (5'-TGGTCATGGATCTGCTAAACC-3').

**Statistical analysis**

Unless otherwise stated, GraphPad Prism v. 10.1.0 was used for all statistical analyses. The number and type of replicates are included in the figure legends. Significance tests were conducted using a one-way ANOVA with a Tukey multiple comparisons test unless otherwise stated. Statistical significance is indicated by asterisks in the figures: * $P \leq 0.05$, ** $P \leq 0.01$, *** $P \leq 0.001$, **** $P \leq 0.0001$.

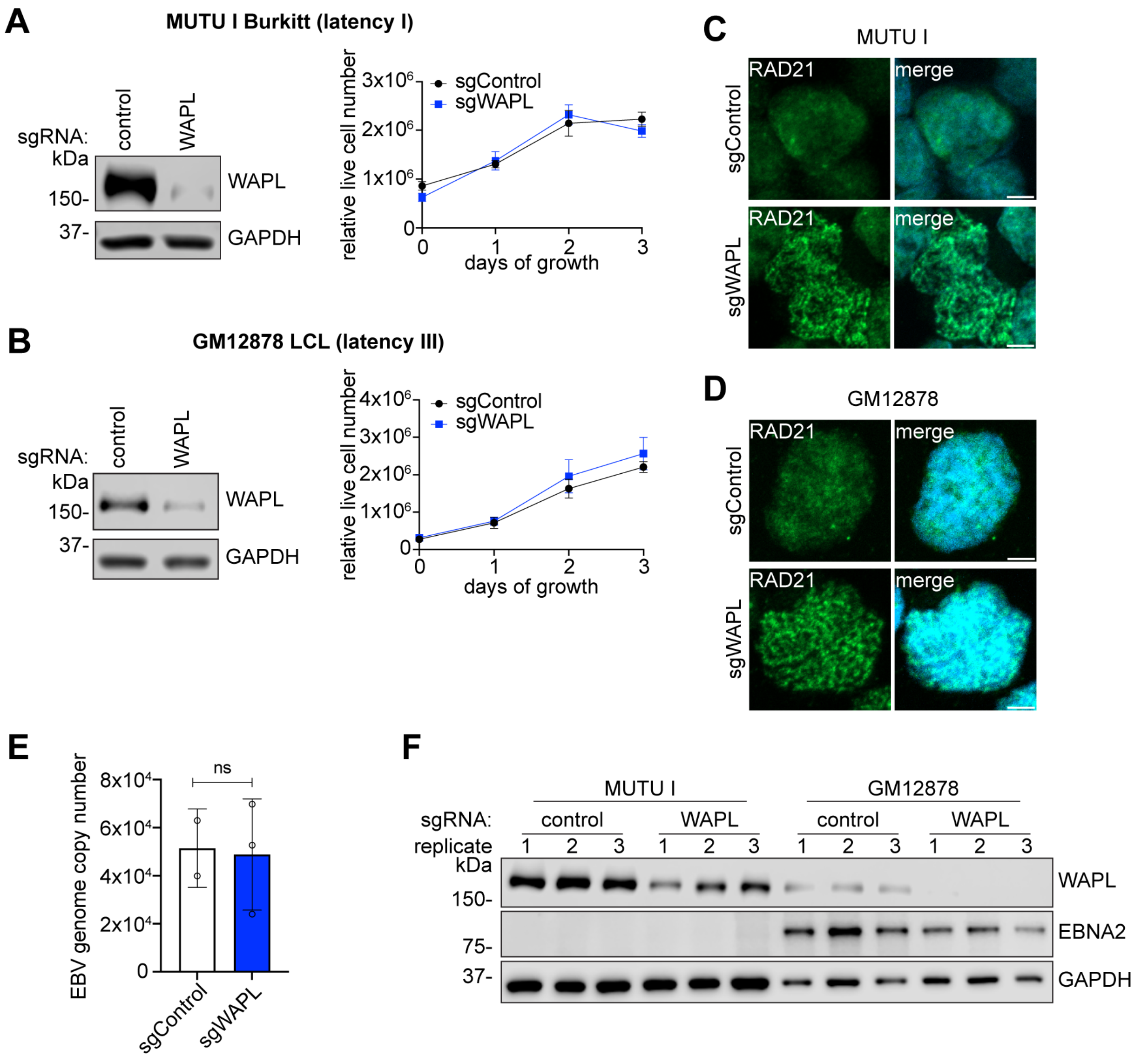


**Figure S1. Burkitt lymphoma cells and LCLs are viable upon CRISPR-mediated WAPL KO.**
**(A, B)** Representative immunoblot analysis (left) demonstrating that sgRNA expression in Cas9+ cells leads to successful WAPL knockout in (A) MUTU I Burkitt lymphoma cells and (B) GM12878 LCLs, and CTG assay (right) indicating that this knockout does not impede cell viability for either cell type. CTG plots show mean relative live cell number ± SD from 3 biological replicates. **(C, D)** Representative immunofluorescence images of RAD21 localization in (C) Cas9+ MUTU I and (D) Cas9+ GM12878 cells expressing control or WAPL sgRNAs. Images are representative of 3 biological replicates. **(E)** Quantification of the number of copies of EBV genomes present in Cas9+ MUTU I cells expressing control or WAPL sgRNAs. Mean ± SD from at least 2 biological replicates. ns = not significant, as calculated by a two-tailed Student's t-test. **(F)** Representative immunoblot analysis of EBNA2 in Cas9+ MUTU I and GM12878 cells expressing control or WAPL sgRNAs. Immunoblots are representative of 3 biological replicates.

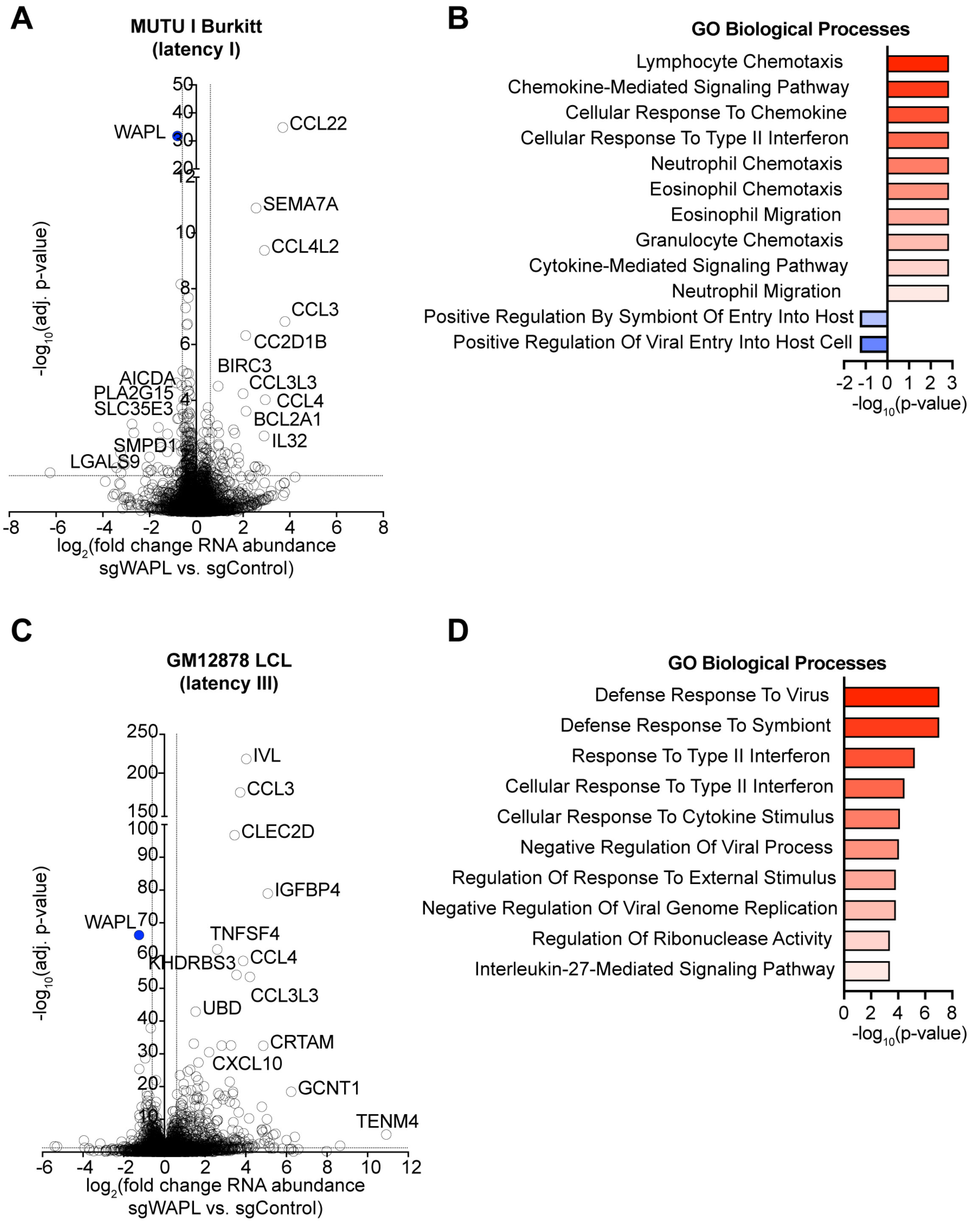


**Figure S2. Burkitt cells and LCLs have similar changes in human gene expression upon CRISPR-mediated WAPL knockout.**
**(A)** Volcano plot of RNA-seq analysis visualizing -$\log_{10}$(adj. p-value) vs. $\log_2$(fold change of human mRNA abundance) from Cas9+ MUTU I Burkitt lymphoma cells and expressing WAPL vs. control sgRNAs, from n = 3 independent biological replicates. **(B)** Significantly altered ($p < 0.05$) GO Biological Processes upon WAPL vs. control sgRNA expression in Cas9+ MUTU I cells. **(C)** Volcano plot of RNA-seq analysis visualizing -$\log_{10}$(adj. p-value) vs. $\log_2$(fold change

of human mRNA abundance) from Cas9+ GM12878 LCLs and expressing WAPL vs. control sgRNAs, from n = 3 independent biological replicates. **(D)** Significantly altered ($p < 0.05$) GO Biological Processes upon WAPL vs. control sgRNA expression in Cas9+ GM12878 LCLs.

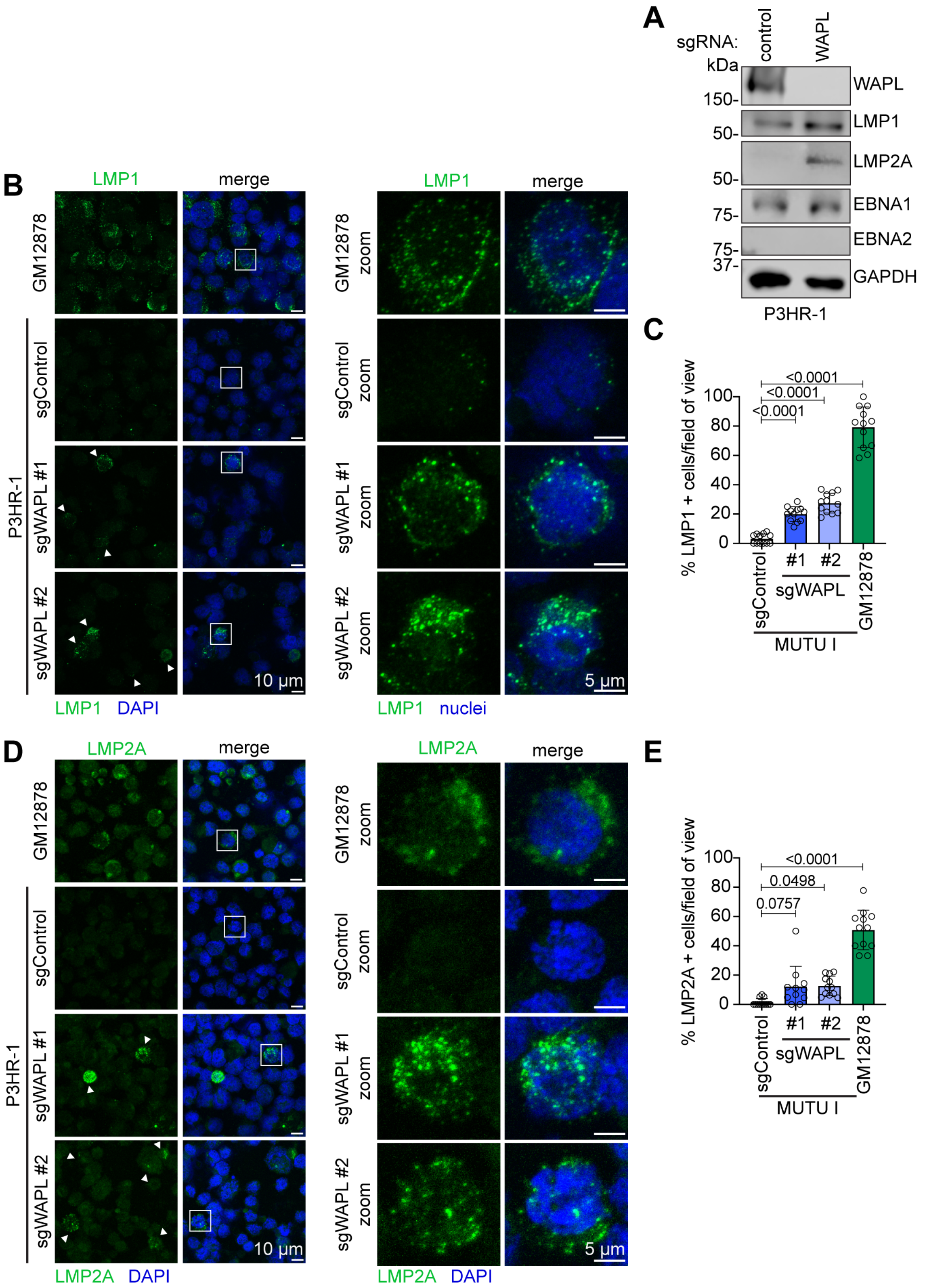


**Figure S3. WAPL loss leads to increased LMP1 and LMP2A levels.**

**(A)** Immunoblot analysis of LMP1 and LMP2A in Cas9+ P3HR-1 Burkitt lymphoma cells expressing WAPL or control sgRNAs. Immunoblot is representative of 3 biological replicates. **(B)** Representative confocal microscopy images from n = 3 biological replicates of anti-LMP1 (green) vs. nuclear DAPI (blue) staining of Cas9+ P3HR-1 cells that expressed control or WAPL sgRNAs, as indicated. Shown at right are zoomed images of a representative cell (indicated by the white box). **(C)** Mean ± standard deviation (SD) percentage of LMP1+ cells per field of view, from n = 3 fields of view from each of three biological replicates. *P*-values shown as calculated by one-way ANOVA. **(D)** Representative confocal microscopy images from n = 3 biological replicates of anti-LMP2A (green) vs. nuclear DAPI (blue) staining of Cas9+ P3HR-1 cells that expressed control or WAPL sgRNAs, as indicated. Shown at right are zoomed images of a representative cell (indicated by the white box). **(E)** Mean ± SD percentage of LMP2A+ cells per field of view, from n = 3 fields of view from each of three biological replicates. *P*-values shown as calculated by one-way ANOVA.

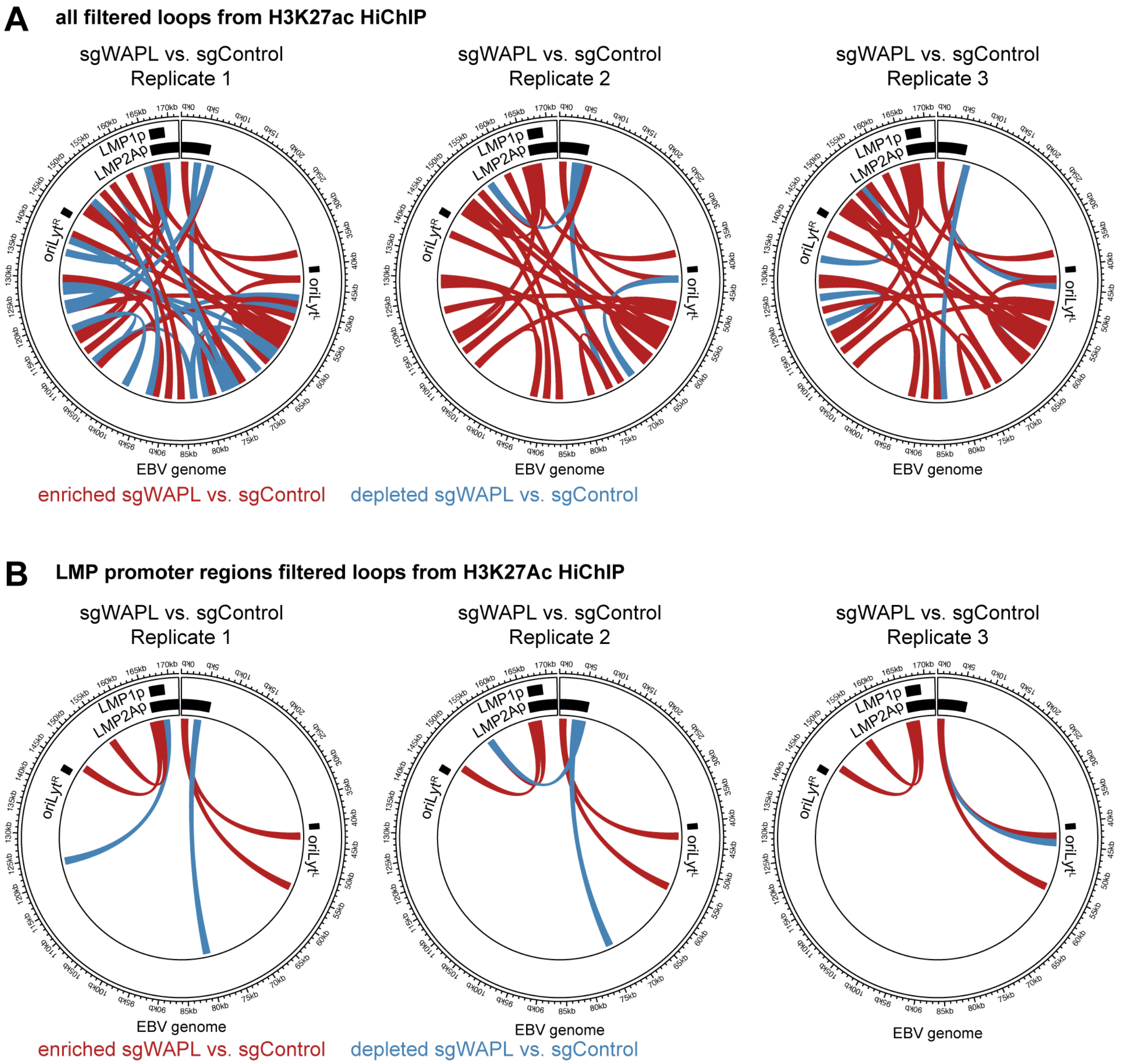


**Figure S4. Loss of WAPL alters looping and enhancer-promoter looping across the EBV genome.**

**(A)** H3K27Ac HiChIP maps of all loops on the EBV genome that are enriched (red) or depleted (blue) in Cas9+ MUTU I cells expressing WAPL compared to those expressing control sgRNAs for each of the 3 biological replicates. **(B)** H3K27Ac HiChIP maps of the LMP promoter loops on the EBV genome that are enriched (red) or depleted (blue) in Cas9+ MUTU I cells expressing WAPL compared to those expressing control sgRNAs for each of the 3 biological replicates.

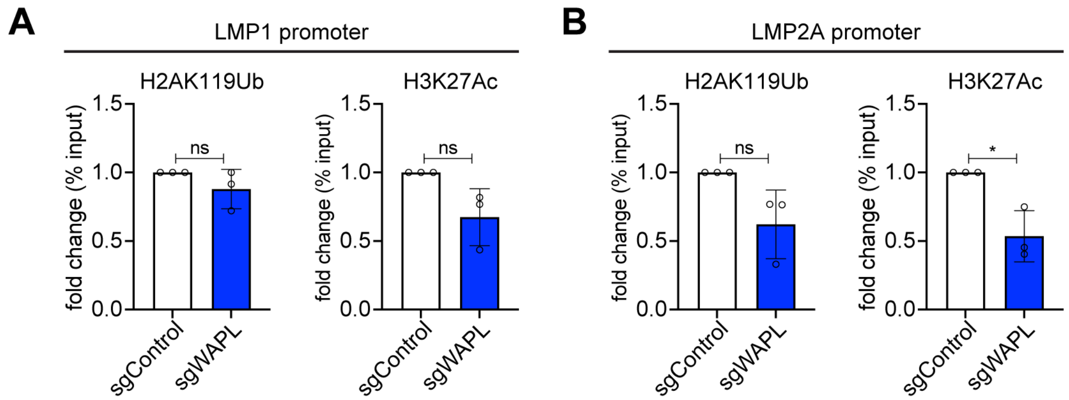


**Figure S5. WAPL knockout does not cause broad changes in epigenetics on the LMP1 and LMP2A promoters. (A, B)** ChIP-qPCR analysis of H2AK119Ub and H3K27Ac accumulation at the EBV (A) LMP1 promoter and (B) LMP2A promoter in Cas9+ MUTU I cells expressing control or WAPL sgRNAs. Shown are mean fold change of the percentage input values ± SD from n = 3 biological replicates. * $P \leq 0.05$, ns = not significant, as calculated by a two-tailed Welch's t-test.
